# Expansion Segment ES30L enriched in birds and mammals can potentially regulate protein synthesis

**DOI:** 10.1101/2022.09.30.510333

**Authors:** Nivedita Hariharan, Sumana Ghosh, Aditi N. Nallan, Arati Ramesh, Deepa Agashe, Dasaradhi Palakodeti

**Affiliations:** Institute for Stem Cell Science and Regenerative Medicine (inStem), Bangalore, India; The University of Trans-disciplinary Health Sciences and Technology, Bangalore, India; National Centre for Biological Sciences (NCBS), Bangalore, India

## Abstract

Ribosomes, the molecular machines that are central to protein synthesis, have gradually been gaining prominence for their regulatory role in translation. Eukaryotic cytosolic ribosomes are typically larger than bacterial ones, partly due to multi-nucleotide insertions at specific conserved positions in the ribosomal RNAs (rRNAs). Such insertions called expansion segments (ESs) are present primarily on the ribosomal surface, with their role in translation and its regulation remaining under-explored. One such ES in the ribosomal large subunit (LSU) is ES30L, which is present only in mammals and birds among eukaryotes. In this study, we show that ES30L possesses complementarity to many protein-coding transcripts in humans and that the complementarity is enriched around the start codon, hinting at a possible role in translation regulation. Further, our in silico analysis analyses and pull-down assays indicate that ES30L may bind to secondary structures in the 5’ UTR of several transcripts and RNA binding proteins (RBPs) that are essential for translation. Thus, we have identified a potential regulatory role for ES30L in translation.

## INTRODUCTION

Eukaryotic cytosolic ribosomes are typically larger than bacterial ribosomes, which is the result of an increase in the content of both ribosomal RNAs (rRNAs) and ribosomal proteins (RPs) (1). For instance, the 25S-28S large subunit (LSU) rRNA is about 3396 nucleotides long in *Saccharomyces cerevisiae* (fungi) and 5034 nucleotides long in *Homo sapiens* (vertebrate), but is only 2904 nucleotides long in *Escherichia coli* (bacteria). The size increase observed in the eukaryotic rRNAs is due to the presence of multi-nucleotide insertions at specific conserved positions, which are termed Expansion segments (ESs) (2–6). Although ESs were first described at least four decades ago, our understanding of their nature and function is still limited.

Some of the main characteristics of these expansions are - 1) Even though the insertion points of ESs coincide, they could be variable in length and nucleotide composition across eukaryotes. For instance, ESs are generally more GC-rich in vertebrates and angiosperms than in invertebrates and single-celled eukaryotes (7). 2) ESs are mainly present on the solvent-accessible surface and do not perturb the core functional centres of the ribosome such as the peptidyl transferase centre, the decoding centre, peptide exit tunnel or the tRNA binding sites (8–11). 3) Initially thought to be nonfunctional insertions, a handful of ESs such as ES7L, ES27L, ES6S and ES9S have been studied and have been shown to play a role in ribosome biogenesis and translation (12–18). With most other ESs remaining unexplored, we have attempted to understand the role of a LSU expansion called ES30L that is specific to endothermic vertebrates (birds and mammals).

In the human ribosome, ES30L is a flexible, tentacle-like extension from helix 78 of the 28S rRNA, which is part of the L1 stalk (19–21). The L1-stalk region from mammalian ribosomes has been shown to interact with IRESs (Internal Ribosome Entry Sites) of some viral mRNAs (22, 23). Given the proximity of ES30L to the L1-stalk and its tentacle-like nature, we have been interested in exploring the potential of human ES30L to interact with mRNAs and proteins. In the current study, using bioinformatic analysis, we show that many protein-coding transcripts possess stretches clustered around the start codon that is complementary to ES30L in *H. sapiens*. Further, our *in vitro* pulldown assays combined with our re-analysis of a published in vivo RNA-RNA interactome dataset indicate that ES30L can potentially interact with many protein-coding transcripts and that such interactions could involve base pairing. We also identify the proteins that ES30L could interact with, using *in vitro* pulldown assays. Together, our results suggest that ES30L can interact with the 5’ UTR of some protein-coding transcripts and also with proteins involved in regulating translation.

## MATERIAL AND METHODS

### Comparison of ES30L among bilaterians

28S rRNA sequences of all the organisms used in this study (85 organisms across 15 phylogenetic groups; refer to Supplementary Table S1A for sequence details) were mined from public databases like Silva LSU r132 (24) and NCBI Nucleotide. Multiple Sequence Alignment (MSA) of the rRNAs was built using Clustal Omega (25) and visualised using Jalview 2.11.2.2 (26). The ESs of the human 28S rRNA were marked in the MSA based on the coordinates taken from Parker et al., (7). The region of the MSA corresponding to ES30L was selected and manually edited in Jalview to improve the alignment. This corrected alignment was used to calculate and plot GC percentage, length and Gap-Adjusted Shannon’s Entropy (GASE) (27) for mammals and birds using custom scripts written in python, bash, awk and R. The ES30L region was also marked in the published human ribosome structure (20) (10.2210/pdb4UG0/pdb) downloaded from PDB. The ribosome structure was rendered using PyMOL (The PyMOL Molecular Graphics System, Version 2.0 Schrödinger, LLC). The secondary structure of LSU rRNA either as a whole or in parts (of the ES30L region) from H. sapiens and the other organisms used in this study was created using the RiboVision web server (28).

### Complementary analysis between ES30L and human protein-coding transcripts

Human protein-coding transcript sequences (83129) were downloaded from GENCODE (v0.29) (29). The human ES30L sequence was extracted from the NCBI reference sequence NR_003287.4 (coordinates: 3975-4035). Ten randomly picked 60 nt stretches from CSL (regions in the 28S rRNA other than the ESs; ES boundaries were taken from Parker et al. (7)) were extracted and also used as a control for the complementarity analysis (Supplementary Table S2-I). The ES30L and the CSL sequences were reverse complemented (only canonical Watson-Crick A:U & G:C antisense pairing considered in this analysis) and used to obtain shorter stretches that range from 7 to 15 nucleotides using the sliding overlapping window approach using custom scripts. For instance, the first 7 nucleotide sequences would span from position 1 to 7 in the reference, the next sequence would span from position 2 to 8 and so on. These shorter stretches from ES30L and CSL were then mapped to the GENCODE transcripts using SeqMap v1.0.13 (30), allowing up to one mismatch in the mappings.

The data presented in this study with mismatch ‘0’ refers to contiguous mapping with zero mismatches. For mismatch ‘1 ‘, data with zero mismatches at lengths of 7 or 8 nucleotides and up to 1 non-terminal mismatch at lengths 9 and above was considered. Only the longest contiguous match (either with 0 or 1 mismatch, as appropriate) from a transcript-ES30L region was considered for further analysis, with the redundant overlapping hits removed using custom awk scripts. Complementary stretches of length 16 nucleotides or more were obtained by extending the overlapping 15 nucleotide matches using a custom awk script. All downstream analyses and plotting were done using custom-written scripts in bash, awk, python and R (31). Gene ontology analysis was done using the shinyGO v0.76 web server (32). Statistical tests used in this study to compare the number of complementary stretches have been referred to from Parker et al. (33). Briefly, Wilcoxon signed-rank test at p<0.05 was used to test the significance of difference in complementarity to transcripts between ES30L and CSL fragments. To characterise the difference in the extent of complementarity from lengths of 7 to 15 or more nucleotides, linear regression was used.

### RNA and Protein pulldown from HEK293T cell lysates with biotinylated RNA oligos

HEK293T cells were cultured in DMEM (Dulbecco’s Modified Eagle Medium) media supplemented with 10% FBS and 1% gentamicin. Cells were grown at 37oC with 5% CO2 for 48 hours and were harvested when they reached 80% confluence. Both the RNA oligos (Supplementary Table S3-I) were synthesized and obtained from Integrated DNA Technologies and included a 5’ biotin tag. The 5’-biotinylated RNA oligos were used in pulldown experiments, following the protocol adapted from Krishna et al. (34). Briefly, 3 μg of the folded biotinylated RNA oligonucleotides in 100 μl of RNA structure buffer (10 mM Tris pH 7.0, 0.1 M KCl, 10 mM MgCl2), was incubated with 1 mg of precleared HEK293T cell extracts in cell lysis buffer (20 mM Tris–HCl pH 7.5, 150 mM NaCl, 1.8 mM MgCl2, 0.5% NP40, protease inhibitor cocktail for mammalian cells (no EDTA), 1 mM DTT, 80 U/ml RNase inhibitor) at room temperature for two hours (100mg of the cell lysate was set aside as input). Post incubation, Dynabeads M-280 Streptavidin™ (11206D, Invitrogen) beads were added to the mixture and incubated for one hour at room temperature. Protein was eluted from the streptavidin beads using SDS buffer (250mM Tris pH 6.8, 10% SDS, 5% beta-mercaptoethanol) followed by acetone precipitation. In case of RNA, TRIzol (Invitrogen)-based extraction from the beads was done. The protein pellets were resuspended in 25mM ammonium bicarbonate and the RNA pellets in nuclease-free water. The total protein/RNA was also extracted from the ‘input’ using either acetone or TRIzol-based methods as appropriate. The isolated protein and RNA samples were sent for mass spectrometry and high-throughput sequencing respectively.

### RNA-sequencing and data analysis

The isolated RNA was analyzed on microfluidic RNA gel electrophoresis with the Agilent 2100 Bioanalyzer before RNA sequencing. Sequencing libraries were prepared using NEBNext® Ultra™ II Directional RNA Library Prep after poly(A) selection from the isolated RNA and sequenced on an Illumina Hiseq 2500 machine (n=2; 2 biological replicates each from ES30L fraction, random fragment fraction and input). Post sequencing, 45 to 64 million single-end (1 * 50 bp) reads were obtained. Adapters were trimmed from the reads using cutadapt v1.8.3 (35) (-a AGATCGGAAGAGCACACGTCTGAACTCCAGTCAC) and were mapped to the GENCODE human transcriptome v0.29 using hisat2 v2.1.0 (36) (default parameters used). Around 80-85% of the reads mapped to the reference transcriptome. The raw read counts mapping to the transcripts were obtained using featurecounts v1.5.0-p1 (37). Normalization of read counts and differential expression analysis was done using the DESeq2 v1.10.1 (38) package in R (31). Transcripts which were two-fold or more upregulated (q<0.05) in the ES30L fraction over the input were considered for further analysis. Plots were generated using the R packages pheatmap (39) and ggplot2 (40). Gene ontology analysis was done using the shinyGO v0.76 webserver (32). Custom bash and awk scripts were used for all downstream analysis.

### LC-MS/MS

The isolated protein was digested using sequencing grade trypsin and the Tryptic peptides were dissolved in 0.1% formic acid (FA) and 2% acetonitrile (ACN). They were then analysed on Thermo EASY-nLC™ 1200 nano System coupled to a Thermo Scientific Orbitrap Fusion Tribrid Mass Spectrometer. Each peptide sample was injected into PepmapTM 100; (75μmx 2cm; Nanoviper C18, 3μm; 100Å) via the auto-sampler of the nano System. Peptides eluted from the peptide trap were separated in a EASY SPRAY PEPMAP RSLC C18 3μm; 50cm x 75μm; 100Å at 45°C. Mobile phase A (0.1% FA in H2O) and mobile phase B (0.1% FA in 80% ACN) were used to establish a 60-min gradient at a flow rate of 250 nl/min. Peptides were then analysed on mass spec with an EASY nanospray source (Thermo Fisher, MA) at an electrospray potential of 1.9 kV. A full MS scan (375– 1,700 m/z range) was acquired at a resolution of 120000, a maximum injection time of 50ms. Dynamic exclusion was set as 20s. MS2 fragmentation was done using HCD technique. Resolution for HCD spectra was set to 30,000 at m/z 100. The AGC setting of full MS scan and MS2 were set as 4E5 and 5E4, respectively. The 10 most intense ions above a 5,000 counts threshold were selected for HCD fragmentation with a maximum injection time of 54 ms. Isolation width of 1.2 was used for MS2. Single and unassigned charged ions were excluded from LC-MS/MS. For HCD, stepped up collision energy (5%) was set to 30%.

### LC-MS/MS data analysis

The raw data (n=4; 2 technical and 2 biological replicates each from ES30L fraction, random fragment fraction and input) was converted to the mascot generic file format using Proteome Discoverer (41) version 1.4 (Thermo Electron, Bremen, Germany) for de-isotoping the LC-MS/MS spectra. The spectral search was performed using a Mascot server (42) (version 2.4.1; Matrix Science, Boston, MA) against the human UniProt database (43) (downloaded on August 10, 2018), with peptide mass tolerance of 10 ppm and fragment mass tolerance of 0.6 Da. Two missed trypsin cleavage sites per peptide were tolerated. Carbamidomethylation (C) was set as a fixed modification, while oxidation (M) and acetylation (N) were variable modifications. Label-free quantification of proteins was based on emPAI values of each identified protein reported by Mascot. All further analysis was done using customised bash, awk and R scripts. The emPAI values were then normalized (44, 45) to compare the relative protein abundance across samples. Only those proteins that were detected in all the four replicates of the ES30L fraction were considered for further analysis and the undetected values from the other samples were replaced with zero. For those proteins that were undetected in all the four replicates in random fragment and input fractions, the value for the first replicate was replaced by 0.000001. Fold change between the pulldown fractions relative to input was computed and Students one-tailed t-test was used to test the significance of fold enrichment, followed by correction for multiple testing using false discovery rate estimation. The proteins that were more than 1.5 fold enriched in the ES30L fraction over input (q<0.05) were probed for gene set enrichment using the shinyGO v0.76 web server (32). The reported ribo-interactome protein list used in this study was taken from Simsek et al. (46).

### Mining and re-analysis of published Splash-Seq and putative IRES datasets

The list of interactions from the in vivo RNA-RNA interactome data (47) was downloaded from GitHub (https://csb5.github.io/splash/). The interactions which included 28S rRNA segments spanning the bin positions 3900 and 4000 (corresponding to ES30L) as one of the interactants were selected for downstream analysis. The reference transcriptome used by the authors was downloaded from their GitHub page and the sequences for interacting transcript regions from the selected interactions were extracted using bedtools v2.25.0 (48). The extent of complementarity to ES30L was analysed in these extracted transcript stretches, using the same pipeline used for the GENCODE (v0.29) (29) transcriptome.

To To check whether any of the transcript regions interacting with ES30L (based on SPLASH-seq data) were part of the putative IRES elements used in this study, putative IRESs from human mRNAs were downloaded from two databases - IRESbase (49) and Human IRES Atlas (50). The sequences for the putative IRES regions were extracted from the transcripts using bedtools v2.25.0 (48) and transcript mapping was done using a nucleotide level blast v2.2.31 (51). The extracted IRES stretches were also checked for complementarity to ES30L using the same pipeline elaborated before. Customised bash and awk scripts were used for data parsing and analysis.

### *In silico* prediction of secondary structural motifs

To probe for the presence of potential structural motifs in the 5’ UTR regions of transcripts that were selectively enriched in the ES30L fraction over input from our RNA pulldown, we considered the 100 transcripts that showed a 50% higher enrichment in ES30L than in the random fragment. Out of the 100, 53 transcripts possessed 229 stretches which were perfectly complementary to ES30L in their 5’ UTR. Sequences that included 20nt flanking these 229 transcript stretches were extracted using bedtools v2.25.0 (48). Any structural motifs present among these sequences were predicted using the covariance model based tool CMfinder v0.2 (52). ATtRACT web server (53) was used to predict binding sites for RBPs on the RNA motifs obtained from the in silico analysis. The secondary structures for the predicted RNA motifs were rendered using the online tool, forna (54). Data parsing was done using custom written bash and awk scripts.

### Data mining and re-analysis of published CLIP-Seq datasets

Published eCLIP-Seq data (fastq.gz files) from K562 cells for RBPs (DHX30, IGF2BP1, NPM1, PCBP1, SRSF1, HNRNPK, U2AF2) were downloaded from ENCODE database (Accession IDs were retrieved from Van Nostrand et al. (55, 56)). For each protein, we considered only the first mate data for two eClip replicates along with one size-matched input for our analysis. The downloaded reads were processed using cutadapt v2.10 (35) to trim adapters and mapped to a 28S rDNA reference (U13369.1:7935-12969) using hisat2 v2.1.0 (36). The read depth per position of the reference was computed using samtools v1.7 (57) and was normalised to per million reads. Fold change (log2 scale) was calculated between the protein pulldown samples and their respective input using custom scripts. In order to compare the binding pattern of PCBP1 between cell types, we downloaded published easyCLIP-Seq data (n=2; size matched input unavailable) from the GEO database (GSE131210) from another study ((55, 56)) done in HCT116 cells. Read depth across 28S rRNA for this dataset was obtained by following the same pipeline used for the other eCLIP-seq dataset. Plots were generated using the R packages pheatmap (39) and ggplot2 (40).

## RESULTS

### A mammal-enriched LSU rRNA expansion segment ‘ES30L’

To investigate the variability in ESs across bilaterians, we sampled 28S rRNA sequences from 85 organisms across various clades (Supplementary Table S1A) and performed a multiple sequence alignment (MSA) using the mined sequences. We marked the boundaries of the LSU ESs on the H. sapiens rRNA and defined the ES regions in the other sequences based on their alignment (Supplementary Figure S1B) to the human sequence. From the MSA, we observed that ES30L was present only in endothermic vertebrates among eukaryotes (Figure 1A). Mapping this region on the structure of the human ribosome showed that ES30L extrudes from the L1-stalk and is usually reconstructed with the help of secondary structure modelling because of its flexibility (Figure 1B, 1C) (19–21). The ES30L stretch was about 3 nucleotides long in rotifers to about 60 nucleotides long in mammals (Figure 1D; Supplementary Table S1A). Amphibians, fishes and arthropods also had shorter expansions in this region, but exhibited a high intra-group variability in length. Further, we also noted that while ES30L was GC-rich in chordates (~92% in *H. sapiens*), it had among the lowest GC-richness in arthropods (Figure 1E; Supplementary Table S1A). To compare the extent of its conservation among mammals and birds, we edited the ES30L region in the MSA manually and used it to calculate Gap-Adjusted Shannon’s Entropy (GASE), which is plotted in Figure 1F. We saw that even though ES30L was well conserved within both clades, the conservation was better among mammals than birds (Figure 1F). Our analysis so far showed that ES30L is a highly GC rich, flexible LSU ES, that is expanded the most and best conserved in mammals. Because of its proximity to the L1-stalk, its tentacle-like nature and its GC-richness, we wanted to investigate its potential to interact with transcripts and proteins.

**Figure 1.**
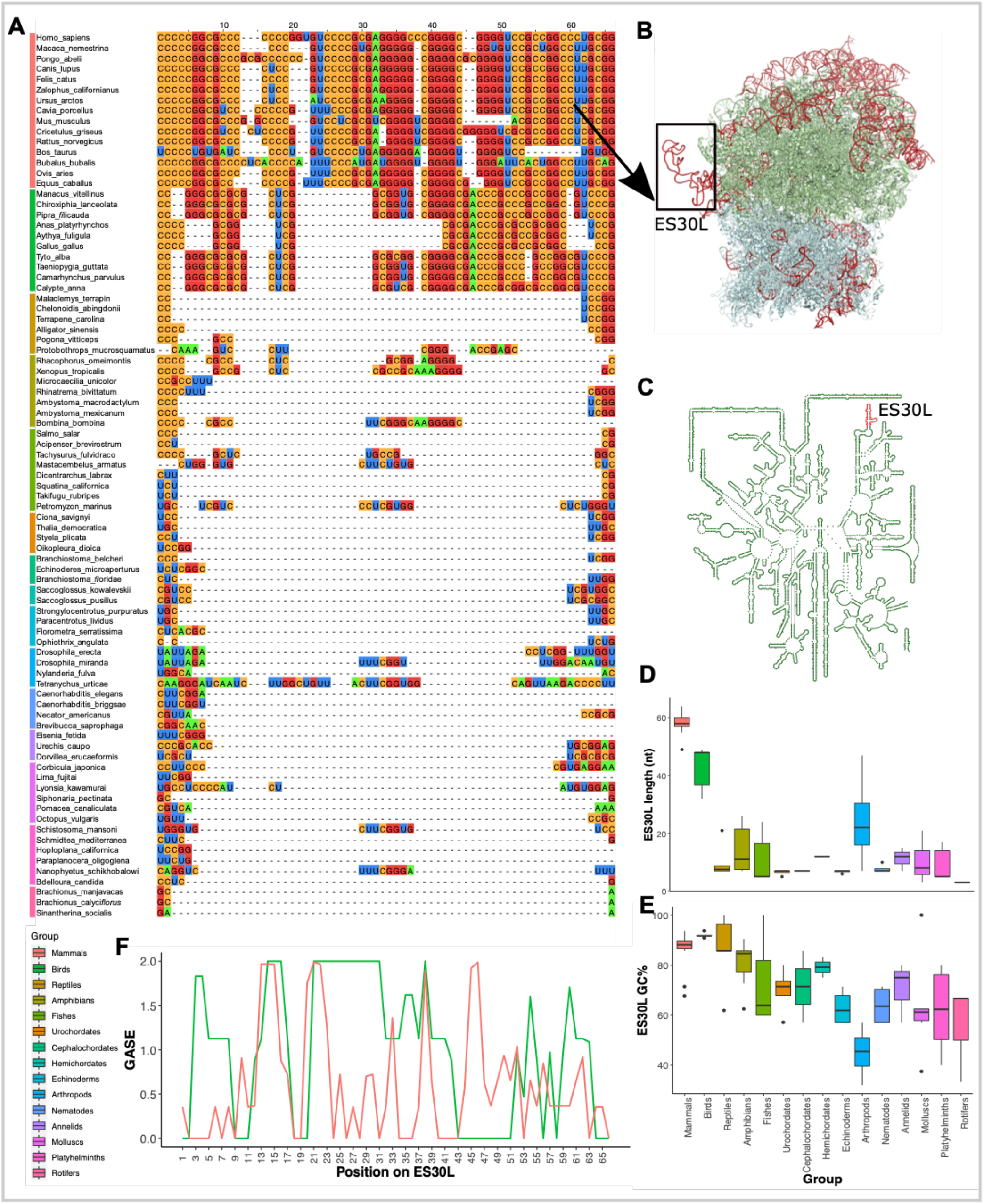
ES30L is a 28S rRNA expansion segment that is expanded largely in mammals and birds. This figure shows the comparison of the ES30L region from different clades (denoted by the coloured bars on the left of the MSA) across bilateria ***(A)*** Multiple Sequence Alignment (MSA) of the ES30L region from 85 organisms across Bilateria. ***(B)*** Front view of the human 80S ribosome (PDB: 4UG0 10.2210/pdb4UG0/pdb) (25). The boxed region on the ribosome is ES30L and corresponds to the rRNA stretch shown in the MSA. It is important to note that the tertiary structure of ES30L is modelled based on both secondary structure prediction and cryo-EM data and is not completely resolved because of its flexibility. The ribosome structure was rendered using PyMOL (The PyMOL Molecular Graphics System, Version 2.0 Schrödinger, LLC). ***(C)*** Secondary structure of the *H. sapiens* 28S rRNA with the ES30L highlighted in red, created using the RiboVision web server (31). The coordinates of ES30L and the other ESs were taken from Parker *et al*., (6). ***(D) & (E)*** Box plots displaying the lengths and the GC% of the ES30L region among the sampled organisms across bilateria. ***(F)*** Gap Adjusted Shannon’s Entropy (GASE) plot for the ES30L region from mammalian (red) and bird (green) sequences shown in the MSA.

### ES30L shows extensive complementarity to many human protein-coding transcripts

To investigate the potential of ES30L to interact with transcripts, we analysed the extent of complementarity between ES30L and human protein-coding transcripts for contiguous stretches starting from a length of 7 nucleotides allowing for zero or one mismatch (see Methods). As a control, regions other than ESs from the 28S rRNA were considered (termed as core segments or CSL here; see Methods; Supplementary Table S2-I) to delineate the specificity of ES30L to the transcripts. Overall, ES30L exhibited a higher degree of perfect complementarity to transcripts than CSL (Figure 2A; 7.25% higher pooled complementary hits with ES30L than CSL; paired Wilcoxon signed-rank test, p < 0.05). A similar trend was observed when one mismatch was allowed, with ES30L possessing 20.85% higher complementarity than CSL (Supplementary Figure S2-IF; p<0.05 in a paired Wilcoxon test). We observed that 84.54% of the transcripts used in the analysis had at least one stretch perfectly complementary to ES30L, with many transcripts having multiple such stretches. This number increases to about 91.77% of transcripts with the inclusion of one mismatch. In case of CSL, nearly 90% of transcripts on an average have at least one complementary stretch, with the proportion going up to 96.57% with the inclusion of one mismatch.

**Figure 2.**
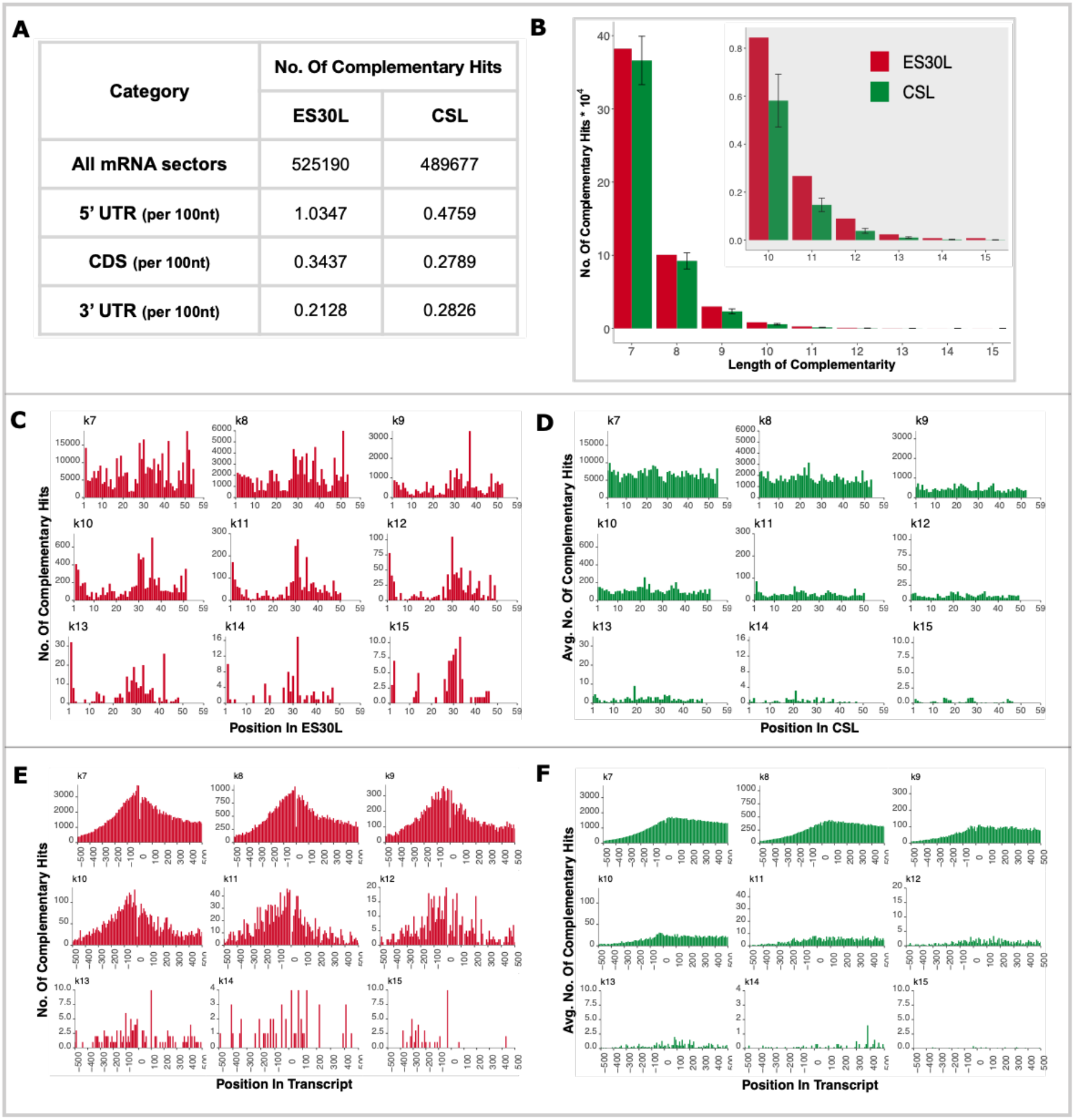
ES30L (from human 28S rRNA) possesses complementarity to many protein-coding human transcripts. This figure shows the summarised comparison between ES30L and CSL (core segment from the LSU; see *Methods*) regions with regards to their contiguous complementarity (zero mismatch) to protein-coding transcripts. In the case of CSL segments, all the data shown here is the average value from the ten segments used in the analysis. ***(A)*** Table showing the total number of complementary hits in all mRNA sectors (5’ UTR, CDS, 3’ UTR) when matched to ES30L and CSL segments. The table also shows the density of complementary hits from each mRNA sector, which is the total complementary hits from that sector divided by the cumulative length of the sector from all the transcripts. ***(B)*** Bar plot showing the number of complementary stretches ranging from a length of 7 to 15 or more nucleotides for ES30L (red) and CSL (green) fragments. The inset in this panel is the data for 10 nucleotides and higher, magnified for better clarity. ***(C)*** & ***(D)*** Histogram showing the number of complementarity stretches to transcripts, arising from each position on ES30L (red) and CSL (green) fragments respectively. ***(E)*** & ***(F)*** Histogram showing the number of complementarity stretches arising from each bin (bin size used here is 10) on the protein-coding transcripts, with ES30L (red) and CSL (green) fragments. The ‘0’ on the x-axis denotes the start codon (TSS) and the negative numbers upstream of it, denoting positions in the 5’ UTR. Only the 500 bases flanking the TSS have been shown in these plots. The complete histogram of this data is provided in Supplementary Figures S2-IC,S2-ID,S2-IIE,S2-IIF.

With 7 and 8 nucleotide stretches, ES30L showed an almost equivalent level of perfect complementarity to the transcripts as CSL, with the extent of pooled hits 5.3% higher in ES30L. With one mismatch, the pooled hits at lengths 7 and 8 were 13.6% lower in ES30L than CSL. More than 50% of the 7 and 8 nucleotide stretches that are complementary (no mismatch) to ES30L got extended to longer stretches (Supplementary Figures S2-IIA, S2-IIB) with the inclusion of one mismatch, with no particular positional preference for the mismatch (Supplementary Figures S2-IIC). But at lengths of 9 nucleotides or higher, the extent of complementarity to the transcripts was 36.12% (34.47% with one mismatch) higher in ES30L than in CSL (Figure 2B; Supplementary Figure S2-IG). Transcript regions complementary to ES30L that are 15 nucleotides or longer were rare, with only 53 such hits (with no mismatch) detected in our analysis. However, we observed 1148 such hits (15 nt and greater) when allowing one mismatch. These 1148 complementary transcript stretches stemmed from 560 genes, whose gene ontology analysis revealed the presence of those that are involved in developmental pathways and neurogenesis among others (Supplementary Figures S2-IH). The presence of such long complementary stretches was much lower in CSL with only 2 hits and 80 such hits on an average containing zero or one mismatch respectively. The slope of the linear regressions on the length of complementary stretch to the log 10 value of the number of complementary hits observed was lower for ES30L than that of CSL (Supplementary Figure S2-IA). The R2 value was equal to or greater than 0.99 in both the cases. The linear regressions were also significant for the data in which one mismatch is allowed. Altogether, our results show that there are more complementary hits of longer lengths in ES30L than in CSL suggesting a greater propensity of ES30L to interact with transcripts.

Along ES30L, around two-thirds of the complementarity to transcripts stemmed from the latter half of the segment (after position 25). This trend was more pronounced at longer complementary stretches, as the percentage of hits from the latter half of ES30L increased from about 62% for 7 nucleotide stretches to about 76% for 15 nucleotide stretches (Figure 2C). This trend was also observed with stretches that included one mismatch (61% for 7 nucleotide stretches to about 72% for 15 nucleotide stretches post the 25th position on ES30L) (Supplementary Figure S2-IID). In case of CSL fragments, the complementarity was present uniformly across the segments, at all lengths (Figure 2D). Even with a mismatch, we observed a similar trend with CSL (Supplementary Figure S2-IIE). Interestingly, the second half of ES30L was well conserved in mammals, with many conserved stretches of guanines (Supplementary Figure S2-IB). The correlation between this sequence conservation in ES30L and the enrichment of complementarity from this region is an interesting aspect that is yet to be explored.

We also analysed the distribution of complementarity to ES30L along the transcripts and observed that the density of such hits was the highest in 5’ UTR when compared to CDS and 3’ UTR. The density of perfectly complementary transcript stretches was about three times higher in 5’ UTR (1.0347 hits per 100nt) than that in both CDS and 3’ UTR (0.3437 and 0.2128 respectively) (Figure 2A). The trend was similar even with the inclusion of a mismatch (Supplementary Figure S2-IF). In the case of CSL, the difference between 5’ UTR and the other transcript sectors was not as stark as that for ES30L, with the density of complementary hits (with zero mismatch) being 0.4759, 0.2789 and 0.2826 for 5’ UTR, CDS and 3’ UTR respectively (Figure 2A). Transcript stretches that are complementary to ES30L, either with or without a mismatch showed the highest enrichment around the start codon (denoted as ‘0’ in the Figures 2E, 2F) and this frequency tapered down with increasing distance from the start codon (Supplementary Figures S2-IC,S2-IIF). We observed a higher enrichment in complementarity (both zero or one mismatch) to ES30L, in the 50 nucleotide stretch upstream of the start codon in the 5’ UTR (4.17% to 4.25% of the total hits) when compared to that observed downstream in the CDS (3.98% or 4.02% of the total hits) (Figure 2E; Supplementary Figure S2-IIF). Complementarity along the transcripts was the highest around the start codon in case of CSL as well (Supplementary Figures S2-ID,S2-IIG), but the enrichment was higher in the 50 nucleotides downstream of the start codon, with 2.33% of hits from CDS as compared to 1.99% from the 5’ UTR (with one mismatch, 2.34% present in CDS versus 1.99% from 5’ UTR) (Figure 2F; Supplementary Figure S2-IIG). These observations hint at a higher potential for the interaction of ES30L with the transcript stretches upstream of the start codon, although interactions with other regions of the transcript is also possible.

Many transcripts had more than one stretch complementary to ES30L. We observed 36 transcripts with over 100 stretches (the highest frequency is 176) and 389 transcripts that possessed between 50 to 100 stretches of varying lengths that are perfectly complementary to ES30L (Supplementary Figure S2-IE). More than 90% of such stretches were either 7 or 8 nucleotides long. With one mismatch, this number was higher (514 transcripts with more than 100 stretches and 3301 transcripts with 50 to 100 stretches). Gene set enrichment analysis showed that the genes that had a hit frequency of 25-50 and 50 or higher, both included genes that are involved in neuronal development and differentiation (Supplementary Figure S2-IE). Our analysis showed that the genes that are involved in neuronal development possess a higher frequency and longer stretches of transcript that are complementary to ES30L, suggesting an increased propensity of such transcripts to bind to ES30L.

### Interactome study reveals binding of ES30L with mRNA potentially mediated by complementarity

To directly test whether ES30L can interact with mRNAs, we used a biotinylated ES30L mimic to pull down RNA from HEK293T cell lysates and sequenced the polyadenylated mRNAs. As a control, we performed a similar pull down with a biotinylated RNA molecule of the same length as ES30L but a random sequence. Both of these experiments were done in duplicates. Our data analysis showed that 1550 protein-coding transcripts were more than two-fold enriched (q<0.05) in the ES30L fraction as compared to the total mRNA (Figure 3A, Supplementary Table S3-II). Out of these, 100 transcripts showed 50% higher enrichment over input in the ES30L fraction when compared to the control fraction (Figure 3B, Supplementary Figure S3-IIIA). Gene ontology analysis of these 100 transcripts showed that this set is enriched in genes that are involved in RNA related processes such as RNA metabolism and translation (Figure 3C). Most of these transcripts (91%) have at least one stretch that is complementary to ES30L with no mismatch (93% transcripts when 1 mismatch is allowed). The number of transcripts with complementarity of varying lengths to ES30L is illustrated in Supplementary Figure S3-IIIB. Most transcripts possessed more than one complementary stretch to ES30L, with 34% of them having more than 10 such stretches.

**Figure 3.**
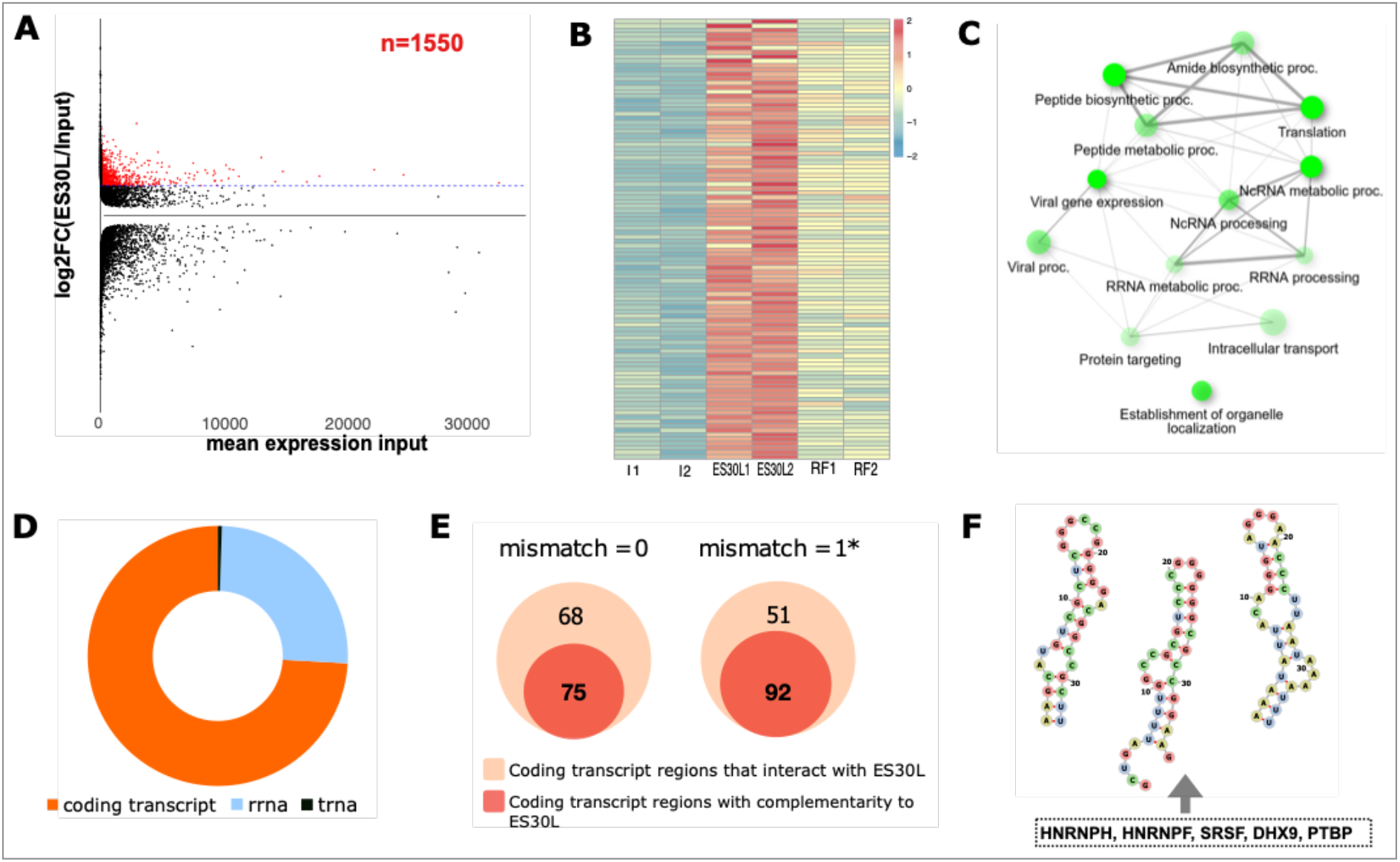
ES30L can potentially interact with protein-coding human transcripts. This figure summarises data that suggests that such an interaction is possible. ***(A)*** MA plot showing the distribution of the fold change of all the transcripts between the ES30L fraction and the input, relative to the mean expression in the input. The points highlighted in red represent the 1550 transcripts that are more than two fold enriched (q<0.05) in the ES30L fraction over the input. ***(B)*** Heatmap showing the expression profiles (normalised read count) of the 100 transcripts that are more than two fold enriched in the ES30L fraction over the input fraction and 50% higher fold change than the random fragment in the pulldown. ***(C)*** Network plot showing the gene categories that are over-represented in the 100 transcripts that are more than two fold enriched in the ES30L fraction over the input fraction and 50% higher fold change than the random fragment in the pulldown. ***(D)*** Pie chart showing the distribution of the RNAs that interact with the ES30L region reported in Aw *et. al. (50). **(E)*** Venn diagram showing the overlap between the number of transcript regions interacting with the ES30L region reported in Aw *et. al. (50)* and the number of those transcript regions harbouring at least one stretch complementary to ES30L. ***(F)*** RNA motifs obtained from *de novo* secondary structure prediction using CMfinder. The sequences used in this prediction were taken from the 100 transcripts from our pull down and were 5’ UTR regions which include stretches complementary to ES30L flanked by 20 nucleotides.

To check whether ES30L-mRNA interactions are possible *in vivo*, we mined a published *in vivo* RNA interactome dataset (47) which has reported pairwise RNA-RNA interactions in HeLa cells, with high sensitivity and specificity using high throughput sequencing based on psoralen crosslinked, ligated, and selected hybrids (SPLASH-seq). From this dataset, we retrieved 193 inter and intra molecular interactions involving the ES30L region of the 28S rRNA (Figure 3D). Out of these, 143 interactions stemmed from protein-coding transcripts, while the rest were mostly with other rRNA regions. We extracted the transcript regions involved in the interactions and checked for presence of complementarity to ES30L in them. Among the 143 interactions, 75 interacting transcript regions contained at least one stretch that is complementary to ES30L and this number increased to 92 with the inclusion of a mismatch (Figure 3E). The 143 interactions involved 118 protein-coding genes, 25 of which were also part of our pull down data. The overlap could possibly vary, since the SPLASH-seq and our pulldown data are obtained from two varied techniques using different cell lines. Out of the 25 genes, 17 had at least one stretch that is perfectly complementary to ES30L (20 genes if one mismatch is included). This data strengthens the possibility that the base-pairing between the complementary stretches could be a way in which the interactions between ES30L and mRNA could be mediated.

Interactions between RNAs could also be mediated by secondary structural elements. To probe this possibility, we mined for predicted IRES (Internal Ribosome Entry Sites) elements (49, 50) in the 118 protein-coding genes from the SPLASH-seq dataset and noted that 14 of them harbour putative IRES (Internal Ribosome Entry Site) sites. These include genes like GAPDH, FTH1, ENO1, RPS11, which are predicted to have IRES in their 5’ UTR. Among the 14 genes, at least 3 (RPS11, FTH1, NHP2) included transcript regions interacting with ES30L (based on reported SPLASH interactions) that are part of predicted IRES (Supplementary Figure S3-IIID). Such regions that could interact with ES30L, also contained multiple stretches that are complementary to it. We then analysed the extent of complementarity to ES30L in all the mined putative IRESs and our data showed that 40.16% of the IRESs have at least one complementary stretch to ES30L (no mismatch), with more than half of them possessing multiple such stretches. Together, our analysis indicated that ES30L can potentially bind to complementary transcript regions, some of which may also be a part of putative IRES structures.

We next wanted to check whether complementary stretches that are present in 5’ UTR of transcripts from our pulldown had any proclivity to form any RNA motifs. Out of the 100 selectively enriched transcripts, 53 had 229 stretches with perfect complementarity to ES30L in their 5’ UTR. We extracted those stretches along with 20 flanking nucleotides and performed an in silico prediction of structural motifs using CMfinder. Our analysis yielded three putative stem-loop motifs that could form in these regions (Figure 3F). The first RNA motif was present in about 12% of the transcripts with the second and third ones present in 4% and 14% of the transcripts respectively. An analysis of putative protein binding sites on these RNA structures using the ATtRACT webserver revealed binding motifs for proteins like HNRNPs, SRSFs, DHX9, PTBP among others. Interestingly, a similar search with ES30L also indicated the presence of binding motifs for the same set of proteins (Supplementary Figure S3-IIIC). This observation prompted us to investigate the protein interactome of ES30L.

### ES30L potentially interacts with several RNA binding proteins

To identify the proteins that potentially interact with ES30L, we did an *in vitro* pull-down in HEK293T cell lysates with a biotinylated ES30L mimic, followed by mass spectrometry (n=4). As a control, we used the same RNA oligonucleotide that was used in our RNA pulldown. We identified fifty-eight proteins that were more than 1.5 fold enriched (q<0.05) in the ES30L fraction when compared to input, which were not enriched in the control fraction (Figure 4A; Supplementary Table S4-I). This set included various categories of RBPs such as: 1)Ribosome biogenesis factors (NKRF, NCL) 2)Helicases (DDX21, DHX30, DHX9) 3)G-quadruplex binding proteins (hnRNPF) 4)Splicing factors (SRSF1, SRSF2, SRSF6, U2AF2) 5)Ribosomal proteins 6)Proteins involved in translation (PCBP1, HNRNPK, IGF2BP1, SRP14, RBM4). Overall, a gene ontology analysis of these proteins showed that many of them are involved in various aspects of RNA metabolism (Figure 4B). Comparison with a published ribo-interactome dataset (46) showed that 42 out of the 58 proteins bind to ribosomes (Supplementary Figure S4-IIA). Interestingly, RNA binding proteins from our data such as the HNRNPs, SRSFs that are also part of the ribo-interactome, have binding motifs in both ES30L and transcript 5’ UTR motifs from our *in silico* analysis discussed earlier (Figure 3F; Supplementary Figure S3-IIC), suggesting that the proteins can bind both RNAs.

**Figure 4.**
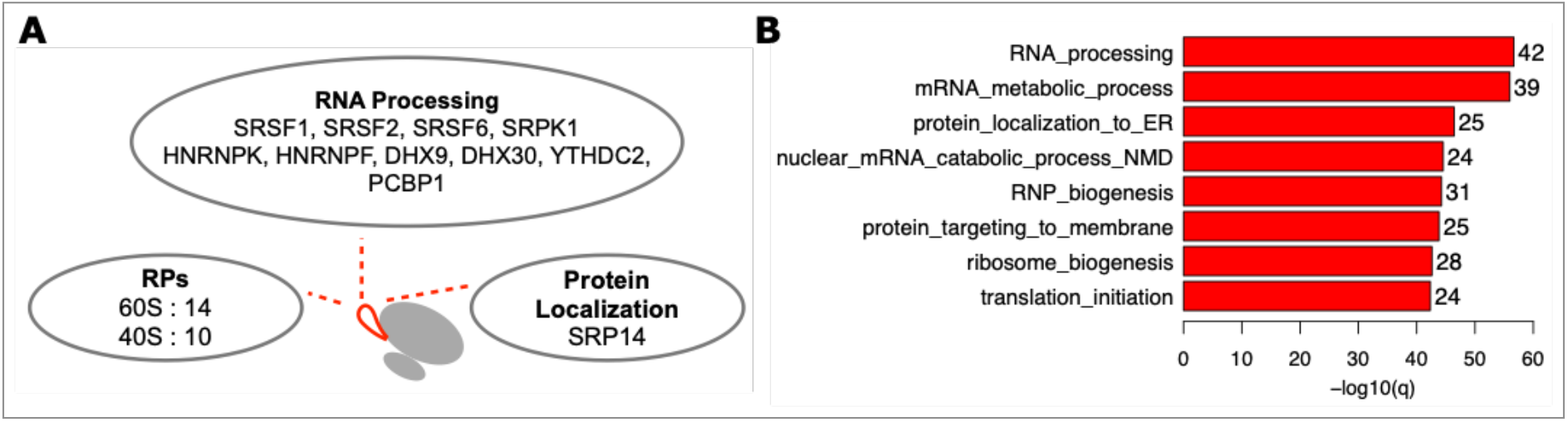
This figure gives an overview of the potential protein interactome of ES30L ***(A)*** Illustration depicting the broad protein categories that were enriched more than two fold (q<0.05) over the total protein fraction in our pulldown. ***(B)*** Bar plot showing the GO (biological process) categories of the proteins that were enriched more than two fold (q<0.05) over the total protein fraction in our pulldown. The details of these proteins are provided in Supplementary Table S4-I.

To probe the binding of ES30L to RBPs *in vivo*, we re-analysed published CLIP-seq datasets (55) from K562 cells for seven proteins that were enriched in the ES30L fraction in our pulldown (Methods; Supplementary Figure S4-IIB). Proteins such as DHX30, NPM1 showed enrichment in reads mapping to ES30L relative to input, suggesting that they could potentially interact with ES30L *in vivo* (Supplementary Figure S4-IIB). The other proteins such as PCBP1, SRSF1, IGF2BP1, U2AF2 or HNRNPK were not enriched in ES30L over input in this dataset. Since *in vivo* molecular interactions could be influenced by their micro-environment, we considered CLIP-Seq data for PCBP1 from two K562 and HCT116 cells. Our analysis of the data from HCT116 cells (55, 56), indicated a preferential mapping of reads towards the latter half of ES30L (coordinates corresponding to nucleotides 4005-4035), with much fewer reads mapping to the first half (Supplementary Figure S4-IID). In contrast, read mapping data from K562 cells (55, 56), showed no such positional mapping preference along ES30L (Supplementary Figure S4-IIC), hinting at the possibility that ES30L:PCBP1 interactions could be context dependent. This needs to be experimentally probed for various RBPs across cell types.

## DISCUSSION

Eukaryotic Eukaryotic ribosomes have accretions both in rRNA and proteins in comparison to the bacterial and archaeal ribosomes, which could have potentially added layers of complexity to the process of translation and its regulation. In our study, we show that ES30L is a highly GC-rich LSU ES that is specific to endothermic vertebrates among eukaryotes. Work by Yokoyama & Suzuki (58) has shown that the ES30L equivalent region (helix 78 of LSU rRNA) in *E. coli* is quite amenable to nucleotide insertions. The ES30L stretch in arthropods (insects) also harbours some expansion, but it is not as well conserved in this group as that in endothermic vertebrates suggesting that this segment could have potentially gained some functional relevance in the latter group. Interestingly, this region is about 30 nt long in the bacteria *E. coli* and *Thermus thermophilus*, 35 nt long in the archaeon *Haloarcula marismortui* while being mostly absent in unicellular and invertebrate eukaryotes (Figure 1A & Supplementary Figure S1D), with longer expansions in most endothermic vertebrates. A detailed phylogenetic analysis may shed some light on whether this rRNA stretch evolved independently in both bacteria and endothermic vertebrates or whether this segment was part of the last common archaeal and eukaryotic ancestor and was lost in unicellular and invertebrate eukaryotes.

Interactions between rRNA and mRNA can occur through base-pairing and a few studies, either experimental or computational (33, 59–63), have explored various aspects of such interactions in both bacteria and eukaryotes. Concrete experimental evidence for an interaction between an ES and mRNA is sparse with a couple of recent studies (17, 64) providing a glimpse into such interactions between ESs and 5’ UTR elements in mRNA. Our bioinformatic analysis shows that ES30L exhibits a high degree of complementarity to many protein-coding transcripts, which is higher than that seen in core rRNA segments. Further, data from our *in vitro pulldown* of transcripts (Supplementary Figure S3-IIB) combined with the *in vivo* RNA-RNA interactome data (Figure 3E) indicate that these transcripts encompass multiple stretches that are complementary to ES30L, hinting at a possibility that such an interaction could involve base pairing. The nature and *in cellulo* impact of these interactions need to be investigated in depth.

The complementarity to ES30L observed across transcripts is denser in the 5’ UTR (Figure 2A; Supplementary Figure S2-IF) with the highest enrichment around the start codon (Figure 2E; Supplementary Figure S2-IC,S2-IIF). Given that ES30L is 92% GC-rich, one reason for the observed trend could be the high 5’ UTR GC content in eukaryotes (65). Data from our study also indicates the presence of complementarity to ES30L in the CDS and 3’ UTR of many transcripts. Our analysis of the *in vivo* RNA-RNA interactome data (47) and data from other studies (66) indicate that *in vivo* interactions between rRNA and mRNA could occur in CDS and 3’ UTR as well.Although the functional relevance of such interactions is yet to be probed, we speculate that they may be involved in RNA metabolic processes such as translational initiation or pausing, mRNA localization and decay.

Given the pervasiveness of the complementarity observed, it is unclear if there would be any selectivity to the potential interactions between the mRNAs and ES30L. We speculate that two factors could contribute to this selectivity: 1)Number/Frequency of complementary stretches present in a transcript, which could increase the possibility of an interaction. 2)The length of the complementary stretches. From our data, we observe that 29.61% of transcripts have at least one stretch complementary to ES30L that is 9 nucleotides or longer (up to 19 nucleotides), which increases further with the inclusion of a mismatch. This raises the possibility that a subset of transcripts could possess even longer stretches with multiple noncontiguous contact points between ES30L and transcript segments. This aspect could be particularly relevant if we factor in the single nucleotide polymorphisms and other variations that could be present in the interacting RNA stretches. Interestingly, we observed that gene sets which possess longer or higher frequency of stretches that are complementary to ES30L, both contain genes that are involved in neuronal development (Supplementary Figure S2-IE,S2-IH). These include genes like SHANK1, SOX11, NTN1, PAX2, CELSR2, ULK1 which have a role in neuron differentiation. Therefore, it would be exciting to probe the functional relevance of ES30L in neurons.

The mere presence of complementarity between two RNA molecules may not ensure an interaction between them, which could also be influenced by the steric availability of both the transcript and ES30L regions possessing complementarity and other *trans*-acting factors. Considering its location on the ribosome, its flexibility and high GC-richness, we think that ES30L possesses a good potential to interact with various RNA Binding Proteins (RBPs). Data from our in vitro protein pulldown shows a selective enrichment of various proteins in the ES30L fraction that are involved in translation regulation and are known to interact with ribosomes (Supplementary Table S4-I; Supplementary Figure S4-IIA). Many of them such as PCBP1, SRSF1, SRSF6, NCL, IGF2BP1, SRP14, DDX21, hnRNPK are known ITAFs (IRES Trans-Acting Factors), which can exert either a stimulatory or an inhibitory influence on IRES-mediated translation. Our bioinformatic analysis also reveals the presence of binding sites for many of these proteins in both ES30L and 5’ UTR stretches from mRNAs enriched in our RNA pulldown (Figure 3F; Supplementary Figure S3-IIC). Such RBPs are known to possess multiple RNA binding domains (67), reinforcing our speculation that they could regulate interactions between ES30 and mRNAs. Concurrent to these observations, our analysis of the SPLASH-seq data also shows that a few of the mRNA stretches that interact with ES30L, are part of putative IRESs (Supplementary Figure S3-IID) and also include multiple stretches complementary to ES30L.

By collating our data with existing literature, we think that ES30L can interact with secondary structural elements in the 5’ UTR, RBPs and influence translation initiation, although such an influence could be either stabilising or inhibitory. Based on our study, we have provided one possible role of ES30L in translation regulation, although it may be involved in other cellular processes too. Since rRNA is distributed as tandem repeats across five chromosomes in the human genome, proving our hypothesis using knock-out experiments is a challenge at present. Given their proposed and established roles in ribosome biogenesis and translation (68), an expansion of the repertoire of experimental techniques is quite important to further our understanding of these enigmatic rRNA segments.

## DATA AVAILABILITY

The RNA-Seq transcriptomic data have been submitted to the NCBI SRA database under the BioProject accession PRJNA883809. The mass spectrometry proteomics data have been deposited to the ProteomeXchange Consortium via the PRIDE(109) partner repository with the dataset identifier PXD036986.

## SUPPLEMENTARY DATA

Supplementary Data are available at NAR online.

## ACKNOWLEDGEMENT

We would like to thank Dr. Sabarinathan Radhakrishnan (NCBS) and Dr. Vinothkumar Kutti Ragunath (NCBS) for their invaluable comments and critical review of the manuscript. We are also thankful for the technical support from the Next Generation Genomics Facility (NGGF) and the Mass Spectrometry Facility at inStem.

