## Supplementary for "Expansion Segment ES30L enriched in birds and mammals can potentially regulate protein synthesis": Supplementary_Figures.pdf

### Supplementary Figure S1B -

The multiple sequence alignment file (in png format) is provided separately.

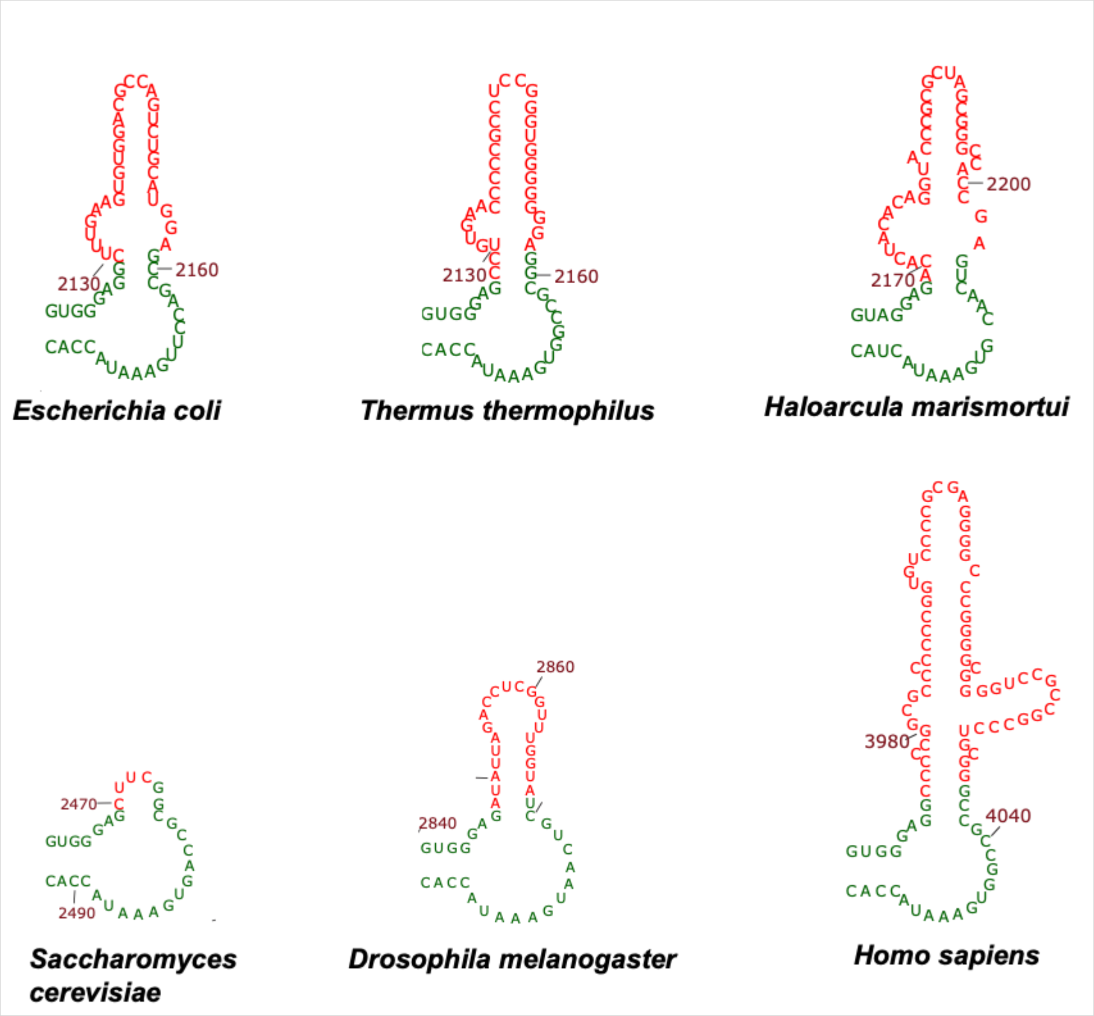


### Supplementary Figure S1D

S1D: Secondary structures of ES30L (red) from different organisms with the flanking core segments of the 28S rRNA shown in green colour. These structures were created using the RiboVision web server.

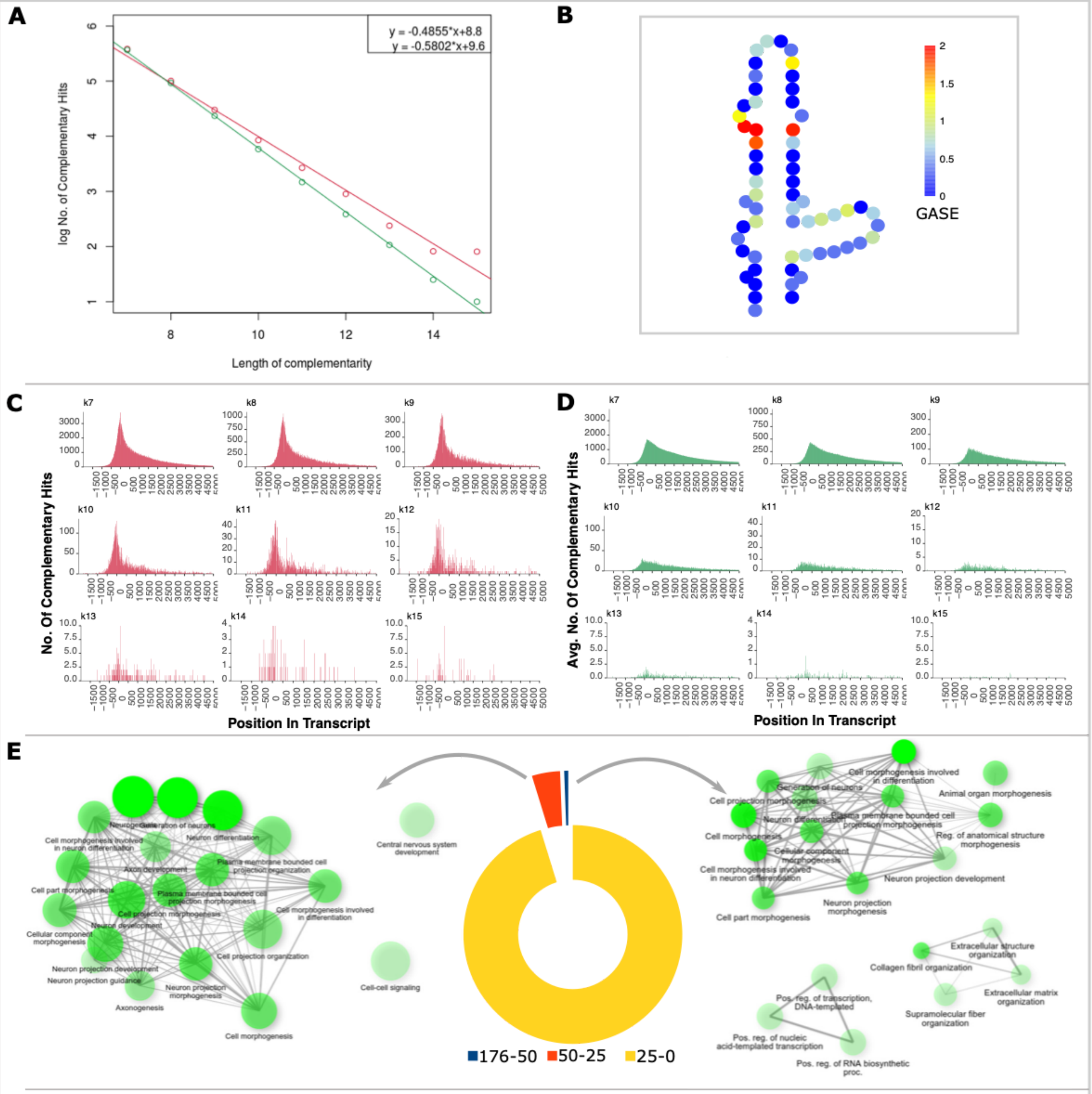


### F mismatch = 1*

| Category | No. Of Complementary Hits | |
|---|---|---|
| | ES30L | CSL |
| All mRNA sectors | 1124509 | 930524 |
| 5' UTR (per 100nt) | 1.9605 | 0.7108 |
| CDS (per 100nt) | 0.6605 | 0.4948 |
| 3' UTR (per 100nt) | 0.3958 | 0.5155 |

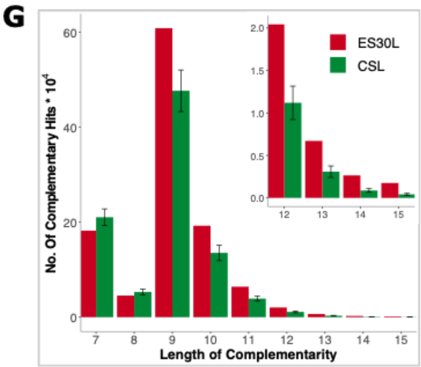


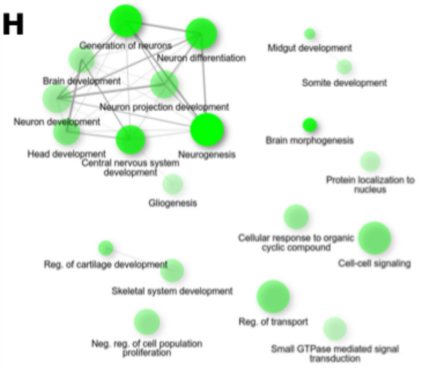


**Supplementary Figure S2-I**

S2-I: ES30L (from human 28S rRNA) possesses complementarity to many protein-coding human transcripts. (A) Linear Regression on the log10 value of the number of complementary hits (with zero mismatch) versus the length of the complementary stretch for ES30L (red) and CSL (green). The inset includes the slopes obtained for the data from both segments (B) Secondary structure representation of the human ES30L with the bases colored according to the GASE values calculated in Figure 1F. This was created using the RiboVision web server (C) & (D) Histogram showing the number of complementarity stretches arising from each bin (bin size used here is 10) on the protein-coding transcripts, with ES30L (red) and CSL (green) fragments. The ‘0’ on the x-axis denotes the start codon (TSS) and the negative numbers upstream of it, denoting positions in the 5’ UTR. (E) Pie chart displaying the proportion of transcripts having varying frequencies of stretches that are complementary to ES30L. The network on the left of the pie chart shows the Gene Ontology - Biological Processes enriched among the transcripts which have 25-50 complementary stretches; while the network on the right shows the processes enriched in the transcripts with more than 50 complementary stretches (F) Table showing the total number of complementary hits in all mRNA sectors (5’ UTR, CDS, 3’ UTR) when matched to ES30L and CSL segments (with up to one mismatch). The table also shows the density of complementary hits from each mRNA sector, which has been calculated by total complementary hits from that sector divided by the cumulative length of the sector from all the transcripts. (G) Bar plot showing the number of complementary stretches ranging from a length of 7 to 15 or more nucleotides for ES30L (red) and CSL (green) fragments. The inset in this panel is the data for 12 nucleotides and higher, magnified for better clarity. In this data, no mismatches were allowed for complementarity at lengths of 7 and 8 nucleotides, with either 0 or 1 mismatch for lengths of 9 nucleotides or higher. (H) Network showing the Gene Ontology - Biological Processes enriched among the transcripts (from 560 genes) that possess stretches of length greater than or equal to 15 nucleotides (with up to one mismatch; see Methods) that are complementary to ES30L.

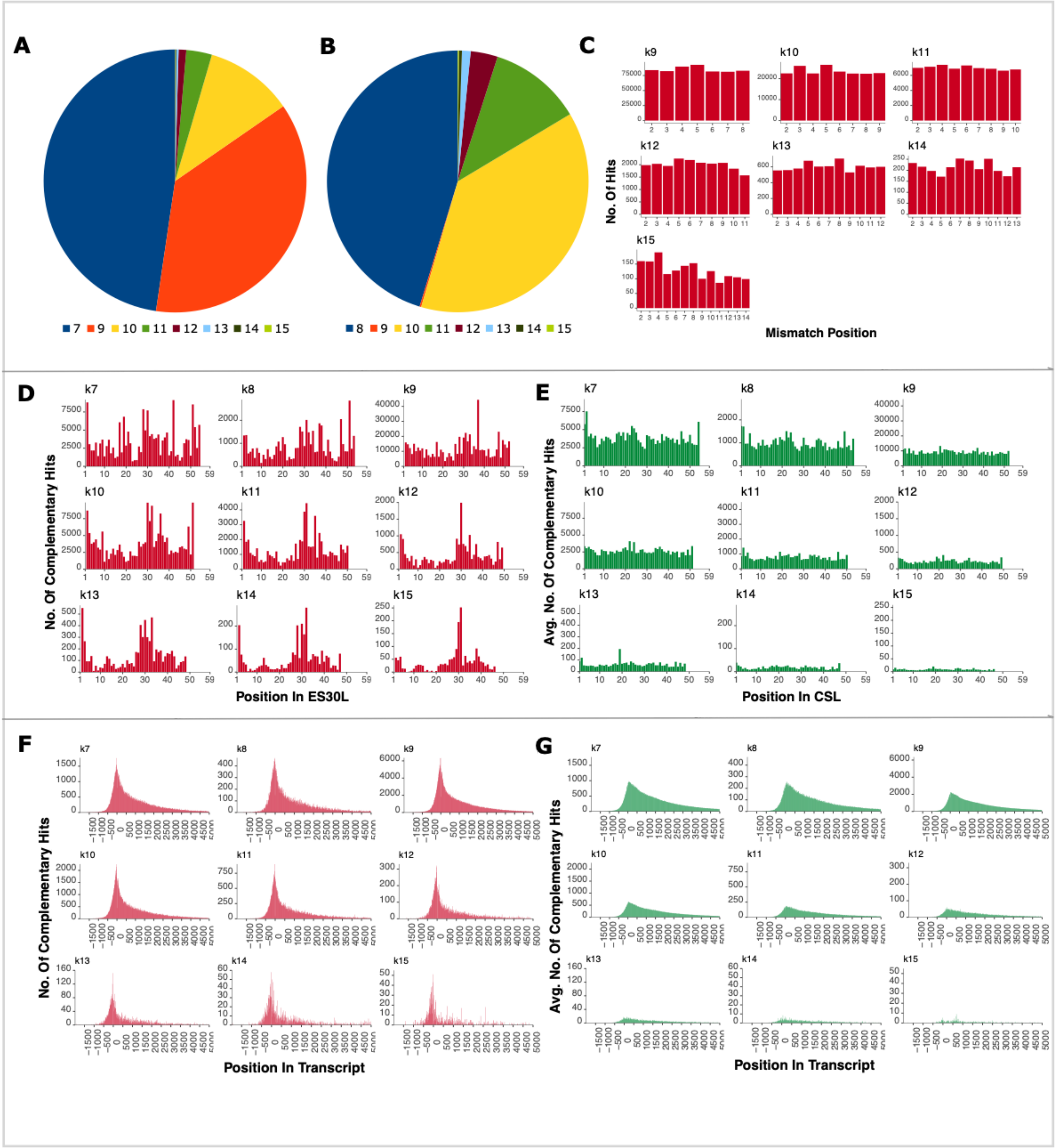


### Supplementary Figure S2-II

S2-II: ES30L (from human 28S rRNA) possesses complementarity to many protein-coding human transcripts. This figure shows the summarised comparison between ES30L and CSL (core segment from the LSU; see Methods) regions with regards to their complementarity (with zero or one mismatch; see Methods) to protein-coding transcripts. In the case of CSL segments, all the data shown here is the average value from the ten segments used in the analysis. (A) & (B) Pie charts showing the redistribution of all the contiguous complementary stretches of lengths 7 and 8 to stretches of different lengths after the inclusion of one mismatch (C) Histogram showing the number of complementary stretches with a mismatch in each non-terminal position within the complementary stretch, at each length. In our analysis, up to one mismatch is allowed for lengths greater than or equal to 9, with none allowed for 7 and 8 nucleotide stretches (Methods). (D) & (E) Histogram showing the number of complementarity stretches to transcripts, arising from each position on ES30L (red) and CSL (green) fragments. (F) & (G) Histogram showing the number of complementarity stretches arising from each bin (bins size used here is 10) on the protein-coding transcripts, with ES30L (red) and CSL (green) fragments. The '0' on the x-axis denotes the start codon and the negative numbers upstream of it, denoting positions in the 5' UTR.

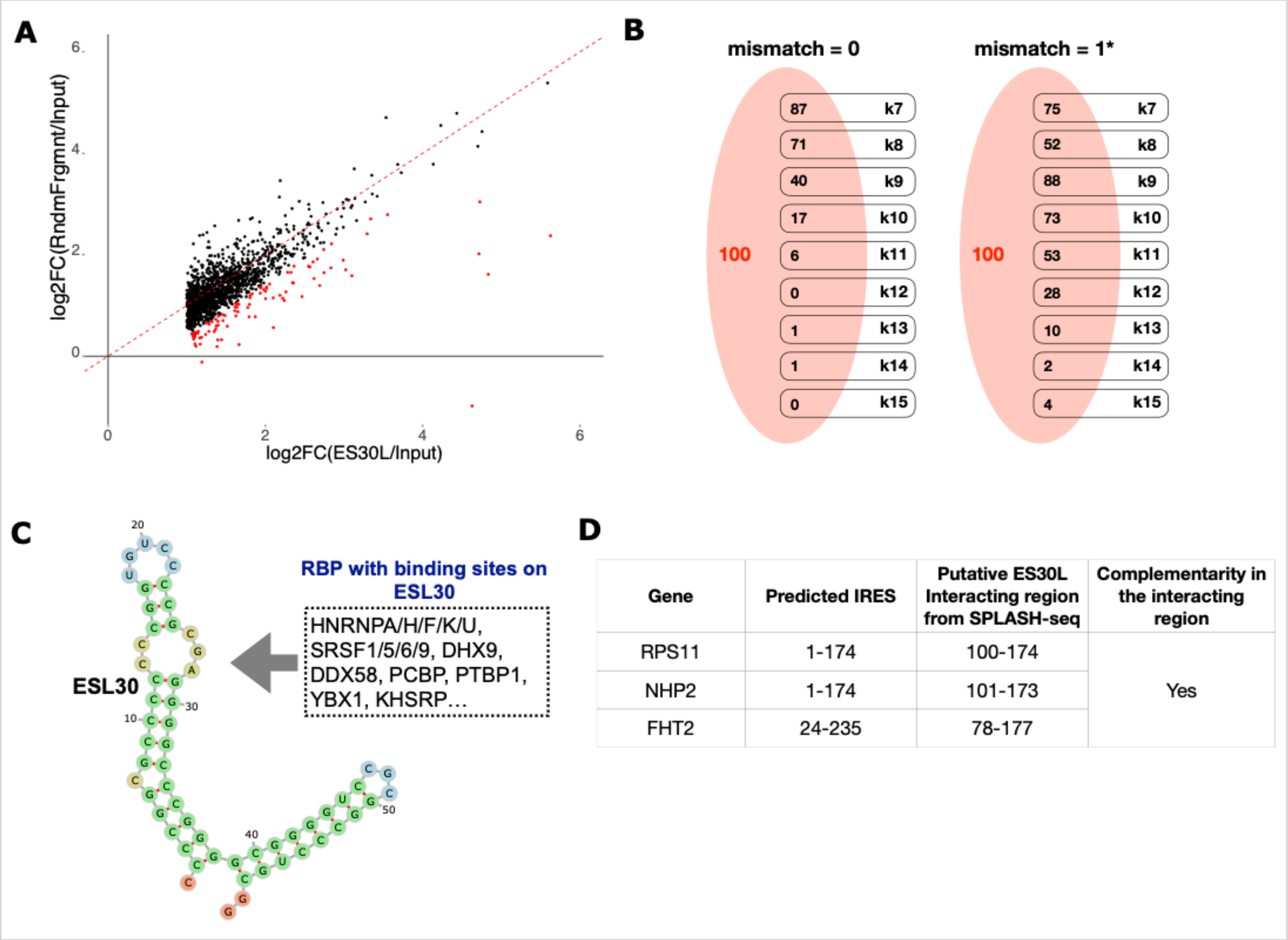


| Gene | Predicted IRES | Putative ES30L Interacting region from SPLASH-seq | Complementarity in the interacting region |
|---|---|---|---|
| RPS11 | 1-174 | 100-174 | Yes |
| NHP2 | 1-174 | 101-173 | |
| FHT2 | 24-235 | 78-177 | |

### Supplementary Figure S3-III

S3-II: ES30L can potentially interact with protein-coding human transcripts. (A) Scatter plot showing the distribution of the 1550 transcripts (Figure 3A) that are more than two-fold enriched in the ES30L fraction over the input and their fold enrichment in the random fragment fraction. Out of the 1550 transcripts, 100 transcripts (coloured in red) showed a 50% higher enrichment over input in ES30L than random fragment (B) Illustration showing the number of transcripts from the 100 pulldown transcripts that have complementary stretches of different lengths with zero and up to one mismatch (see Methods) (C) Predicted secondary structure of ES30L along with the putative RNA binding protein motifs that are present on it. (D) Table enlisting the ES30L interacting regions from three genes that are also part of putative IRESs.

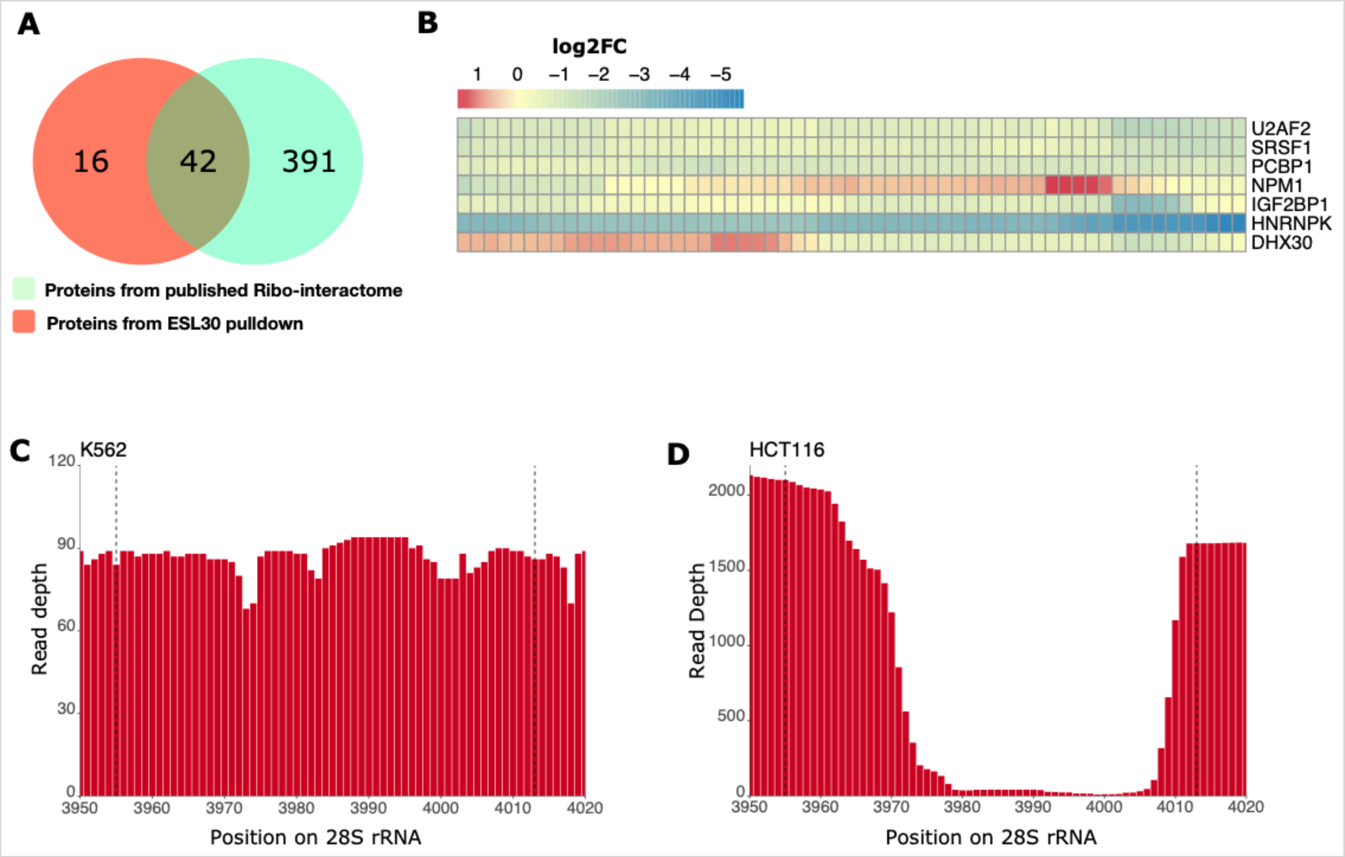


### Supplementary Figure S4-II

S4-II: ES30L could interact with RBPs. (A) Venn diagram showing the overlap between the proteins identified in our pulldown experiment and a published ribo-interactome (Shi et al. 2017). (B) Heatmap showing the enrichment of reads (for the protein fraction over their size-matched input) in the ES30L region from published CLIP-seq data (Van Nostrand et al. 2020) for eight proteins that were identified in our pulldown. (C) & (D) Coverage plot showing the read depth pattern across the ES30L stretch (U13369:3955-4013; Supplementary Table S1C) from two different studies - Van Nostrand et al. 2020 (left; K562 cell line) and Porter et al. 2021 (right; HCT116 cell line).
