## Supplementary figures and images for "Expansion Segment ES30L enriched in birds and mammals can potentially regulate protein synthesis"

### Supplementary_Figure_S1B.png

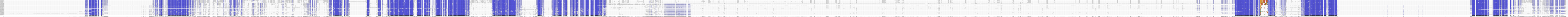
